# Dynamic Changes in Fecal Bacterial Microbiota of Dairy Cattle across the Production Line

**DOI:** 10.1101/2020.02.21.960500

**Authors:** Lei Zhao, Xunde Li, Edward R. Atwill, Sharif Aly, Deniece R. Williams, Paul Rossitto, Zhengchang Su

**Affiliations:** Department of Bioinformatics, the University of North Carolina at Charlotte, Charlotte, North Carolina, USA; Western Institute for Food Safety and Security, University of California, Davis, California, USA; Department of Population Health and Reproduction, University of California, Davis, California, USA; Veterinary Medicine Teaching and Research Center, University of California, Davis, California, USA

**Keywords:** 16S rRNA, dairy cattle, microbiota, feces, GIT, NGS, bioinformatics

## Abstract

Bacteria play important roles in the gastrointestinal tract (GIT) of dairy cattle as the communities are responsible for host health, growth and production performance. However, a systematic characterization and comparison of microbial communities in the GIT of cattle housed in different management units on a modern dairy are still lacking. In this study, we used 16S rRNA sequencing to evaluate the fecal bacterial communities of 90 dairy cattle raised and housed in 12 clearly defined management units on a modern dairy farm. We found that α-diversity differed between several pairs of management units, especially between the hutch calves (management unit 1) and the other units except post weaned heifers (management unit 2). Although β-diversity revealed that most of the samples did not cluster according to their management unit membership except management unit 1, in which the samples grouped and separated from others, we observed three major clusters. Besides the hutch calves (management unit 1), most samples from the other 11 units formed two distinct clusters and the relative abundance of Patescibacteria might be the reason for the separation. Moreover, we identified 212 amplicon sequence variants (ASVs) with relative abundances > 0.1% in more than 10% of the total samples that had significantly different abundances in these units. Furthermore, we found that, as the calves aged, 19 ASVs that were exclusively detected in unit 1, gradually degraded and never reoccurred in other management units. Lastly, we recognized 181 pairs of interactions between 61 ASVs with possibly strong synergistic or antagonistic relationships. These results suggest the enteric microbial communities of dairy cattle housed in different management units are quite dynamic.

**IMPORTANCE:** Bacterial communities of GIT are crucial for ruminants, such as dairy cattle since they contribute to not only the cattle’s physiology and health but also milk production and food safety that are closely related to human health. Both scientifically and agriculturally, it is necessary to have deep insights into the composition and changes of the bacteria on modern dairy farms. In this study, we provided the profiles of fecal bacterial communities of dairy cattle in each management unit on a modern dairy and described how the enteric microbial communities changed across these management units. The findings have implications for improving animal health, dairy production, farm management, and food safety.

---

It is now well-established that microbiota, especially bacterial communities, play crucial roles in the physiology and health of all mammals. While the number of bacterial cells in the human body has been estimated as 10 times more than the number of human cells (1) or at roughly the same order in a recent study (2), this ratio can rise to approximately 120 times in ruminants such as cattle (3). Cattle depend on their gastrointestinal microbiota to digest and convert the plant mass that cannot be directly digested into absorbable nutrients necessary for host health and development. Thus, a better understanding of the structure of the gastrointestinal microbiota is instrumental for both production and scientific inquiry. Particularly, in the modern dairy system, calves, heifers and cows at different ages and production status (Table 1) are raised and managed in independent yet inter-connected managements units (Fig. 1), so it is important to understand the dynamics of the gut microbiota changes between/among different management units over the entire production life cycle of dairy cattle. The knowledge gained can help design new strategies to improve production as well as the health of both the animals and humans who consume the produced meat and milk.

**TABLE 1.** Classification of cattle based on the growth, production stages, and their residential management units of the dairy herd where samples were collected.

| **Management Unit** | **No. of Samples** | **Unit ID** | **Description** |
| --- | --- | --- | --- |
| Hutch calves | 9 | 1 | From birth to approximately 1-2 weeks after weaning, individually housed (1 to 70 days age approximately) |
| Post weaned heifers | 6 | 2 | Group-housed heifers and bull calves fed solid diet |
| Breeding heifers | 8 | 3 | Approximately 13 to 15 months old |
| Springer | 8 | 4 | Within 1 to 4 weeks of calving, pregnant nulliparous heifers |
| Fresh uniparous cows | 8 | 5 | 1 to 2 months post-calving, first lactation cows |
| Mid-lactation uniparous cows | 8 | 6 | 60 to 250 Days in Milk, first lactation |
| Fresh multiparous cows | 8 | 7 | 1 to 2 months post-calving, second lactation or greater |
| Mid-lactation multiparous cows | 8 | 8 | 60 to 250 Days in Milk, second lactation or greater |
| Pregnant late lactation cows | 8 | 9 | >250 Days in Milk |
| Far-off dry cows | 3 | 10 | 21 to 60 days prior to calving, multiparous dry cows |
| Close up dry cows | 8 | 11 | within 21 days prior to calving, multiparous dry cows |
| Hospital pen | 8 | 12 | Lactating cows treated with medication that requires milk withdrawal |

Since traditional culture-dependent techniques can only recover a small portion of the microbial population (4), several research groups have probed the gut microbiota of dairy cattle using the next-generation sequencing (NGS) technologies (3, 5–12). For instance, Mao et al. analyzed the microbiota of ten gastrointestinal sites in Holstein cattle using 16S rRNA sequencing and found that these cattle hosted microbiota with significant spatial heterogeneity (6). Dill-McFarland et al. analyzed the succession pattern of bacterial communities in dairy cattle from 2-week-old to first lactation (8). Shanks et al. profiled the structure of fecal bacteria in cattle from various feeding operations (11). However, a systematic contrast of the gut bacterial microbiota in dairy cattle along all the management units on a modern farm is still absent. To fill these gaps, we analyzed the bacterial communities of 90 fecal samples of dairy cattle sampled from 12 management units of a dairy herd in California starting from the newborns to late lactation and dry cows using 16S rRNA sequencing data (Table 1 and Fig. 1).

**FIG 1.**
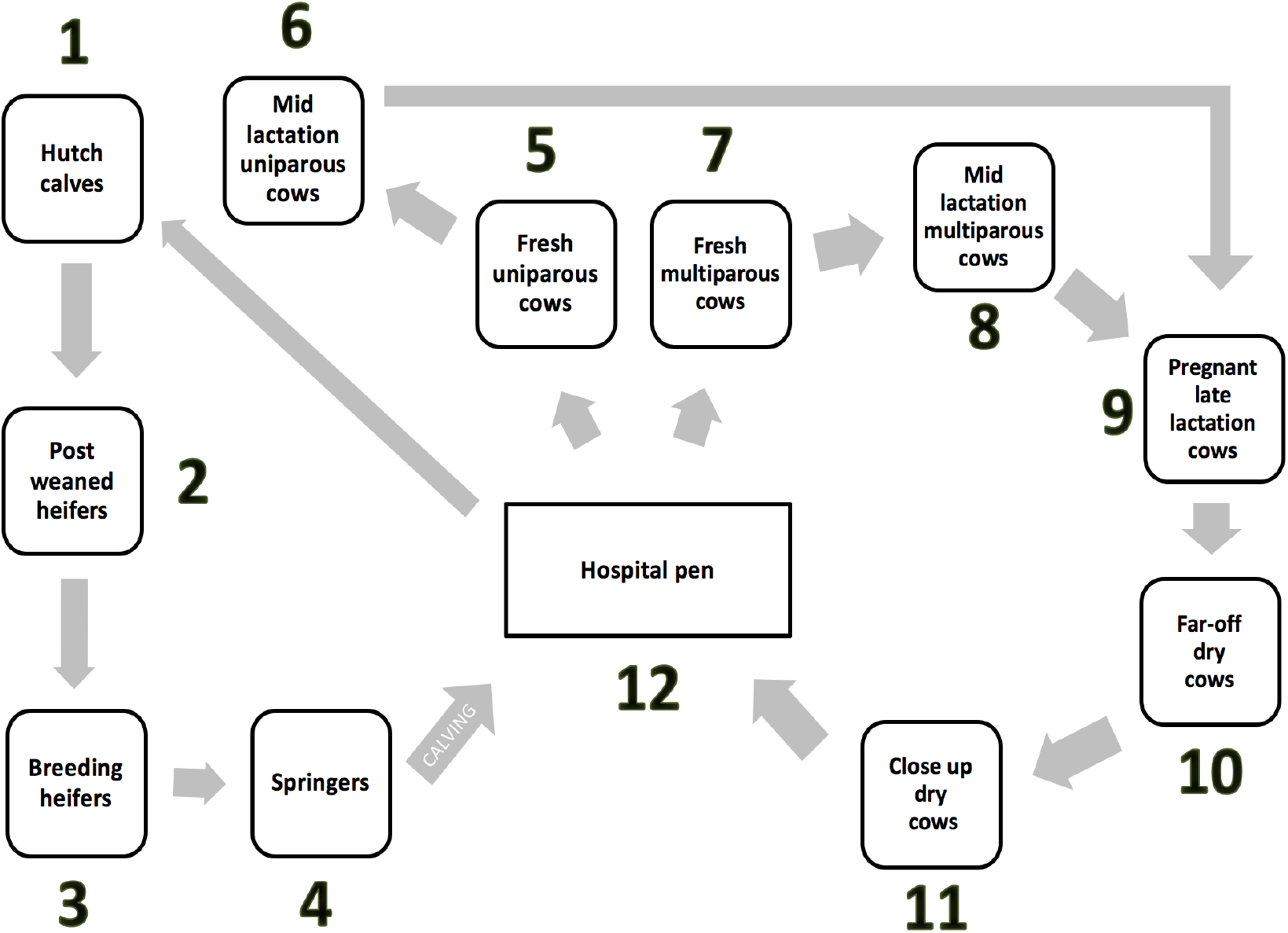
Schematic diagram of the independent yet inter-connected management units of the dairy herd where the samples were taken. The number around each box represents the management unit ID used in this study.

## RESULTS

### Composition of the bacterial communities in the fecal samples

We generated a total of 6,092,309 16S rRNA paired-end reads with an average of 67,692 ± 7,963 reads per sample. Clustering sequence reads from V3/V4 regions of 16S rRNA genes into different Operational Taxonomy Units (OTUs) has been a widely adopted strategy. However, the traditional OTU methods fail to tell the technical errors from real biological variations (13). Thus, we used a new method DADA2 that infers sequence variants by learning the sequencing error rate from the samples (13), and adopted the term “Amplicon Sequence Variants (ASVs)” instead of the traditionally used term OTUs (14) to refer to unique V3/V4 regions of 16S rRNA genes in the samples. After quality filtering and chimera removal with DADA2 (13), a total of 5,080,044 sequence reads with an average of 56,444 ± 6,633 reads per sample were preserved. ASVs taxonomically classified as “Archaea” and “Eukaryote” were excluded and ASVs that appeared in less than 9 samples (10% of sample size) were also excluded. The ASV table was normalized by rarefying with a threshold of 12,637 sequences (see Materials and Methods), which is the minimum number of reads of all the 90 samples after processing. All downstream analyses were based on the normalized ASV table (supplemental file 1: normalized ASV table).

We identified a total of 4,681 ASVs in the 90 fecal samples. These 4,681 ASVs were detected in varying number of samples, and only three ASVs classified as Candidatus_saccharimonas (ASV-1, ASV ID can be found in supplemental file 2), Ruminococcaceae_UCG-014 (ASV-11), and [f]Lachnospiraceae (ASV-15), were detected in every sample (Fig. S1A in the supplemental file 2). Each sample had a different number of ASVs (Fig. S1B in the supplemental file 2) and the ASV relative abundances were variable between samples (Fig. S1C in the supplemental file 2). Out of the 4,681 ASVs, 4,486 (95.83%) had an average relative abundance < 0.1%, and 195 (4.17%) ASVs had relative abundance ≥ 0.1%. The ASVs having an average relative abundance ≥ 0.1% accounted for 599,769 (52.73%) reads of the total reads.

The 4,681 ASVs could be taxonomically assigned to 20 phyla, of which Firmicutes (2,573/54.97%), Bacteroidetes (1,291/27.58%) and Tenericutes (267/5.70%) included the highest number of the ASVs (Fig. 2A). While only Firmicutes (587,075/51.62% reads), Bacteroidetes (412,852/36.30% reads), Patescibacteria (53,402/4.70% reads) and Proteobacteria (16,379/1.44% reads) were observed in all 90 samples, they accounted for 94.05% of the total bacterial communities at the phylum level. Firmicutes had the highest average relative abundance in the samples from all the management units except unit 1, where Bacteroidetes accounted for 62.25% and Firmicutes for 33.37% at the phylum, respectively (Fig. 2B).

**FIG 2.**
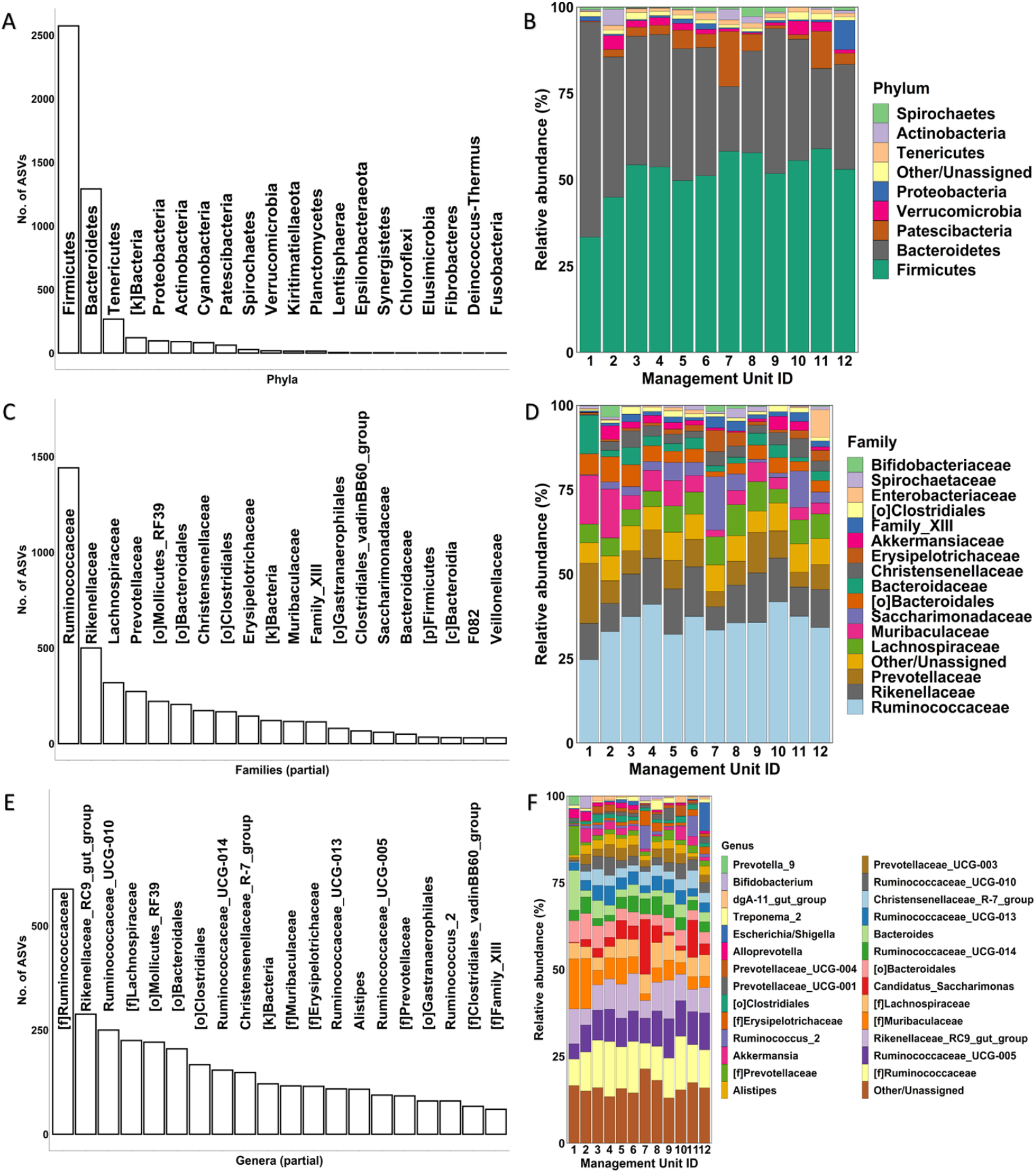
Assignment of the ASVs to different taxonomic levels. (A) The number of ASVs assigned to different phyla. (B) Average relative abundances of the phyla in the samples from different management units. (C) The number of ASVs assigned to different families. (D) Average relative abundances of the families in the samples from different management units. (E) The number of ASVs assigned to different genera. (F) Average relative abundances of the genera in the samples from different management units. In B, D, and F, the length of color-coded bars represents the average relative abundance of the taxa in the samples in the indicated management units. Taxa with < 2% relative abundance in all the units were merged into the “Other/Unassigned” category.

At the family level, the 4,681 ASVs could be assigned to 99 families, in which Ruminococcaceae (1,441/34.91%), Rikenellaceae (500/11.58%), Lachnospiraceae (319/6.78%), and Prevotellaceae (273/8.47%) contained the highest number of the ASVs (Fig. 2C). Although only 12 families were detected in all 90 samples, they accounted for 992,097 (87.23%) reads of the total bacterial communities at this taxonomic level. Ruminococcaceae (397,075/34.91% reads) was the most dominant taxon in average relative abundance at this level, followed by Rikenellaceae (131,673/11.58% reads), Prevotellaceae (96,322/8.47% reads), Lachnospiraceae (77,143/6.78% reads), Muribaculaceae (70,454/6.19% reads) and Saccharimonadaceae (53,121/4.67% reads) (Fig. 2D).

At the genus level, although 2,169 (46.34%) of the 4,681 ASVs could be assigned to 167 known genera, the remaining 2,512 ASVs (53.66%) could not be classified to known genera. These unclassified taxa might be novel bacteria in the cattle feces or not differentiable solely based on the hypervariable regions of 16S rRNA genes. Overall, the largest number of the ASVs were assigned to [f]Ruminococcaceae (588/11.92% ASVs), followed by Rikenellaceae_RC9_gut_group (288/6.15% ASVs), and Ruminococcaceae_UCG-010 (250/5.34% ASVs) (Fig. 2E). While only 17 genera were consistently detected in the 90 samples, these commonly shared genera occupied 72.81% of the total bacterial communities at this taxonomic level. Most of the assigned genera had an average abundance of < 2% in the samples from the management units, while only 15 genera had a relative abundance ≥ 2% including [f]Ruminococcaceae (11.52%), Ruminococcaceae_UCG-005 (8.75%), Rikenellaceae_RC9_gut_group (8.20%), [f]Muribaculaceae (6.19%), [f]Lachnospiraceae (5.16%), Candidatus_Saccharimonas (4.67%), [o]Bacteroidales (4.25%), Bacteroides (3.57%), Ruminococcaceae_UCG-014 (3.48%), Ruminococcaceae_UCG-013 (3.40%), Christensenellaceae_R-7_group (2.97%), Ruminococcaceae_UCG-010 (2.61%), Prevotellaceae_UCG-003 (2.50%), [f]Prevotellaceae (2.23%), and Alistipes (2.17%) (Fig. 2F).

### Alpha diversity of the cattle fecal bacterial communities in the management units

We next compared the richness and evenness of the fecal bacterial communities of cattle in different management units in the production line using both Chao 1 richness index and Shannon diversity index. As shown in Figs. 3A and 3B, the fecal bacterial communities in pre-weaned calves fed primarily milk and housed in individual hutches (unit 1) had significantly lower Chao 1 and Shannon indexes than those in all other units, except for the post-weaned group-housed calves fed a solid diet (unit 2) (Kruskal Wallis, false discovery rate, FDR < 0.05), suggesting that the hutch calves generally had simpler fecal bacterial communities. Post weaned heifers (unit 2), while having no significant differences in bacterial communities compared with hutch calves (unit 1), consistently differed from breeding heifers (unit 3), springers (unit 4), and pregnant late lactation cows (unit 9), which may be related to the development of the rumen and these growing young cattle being full ruminants by the time they are bred.

**FIG 3.**
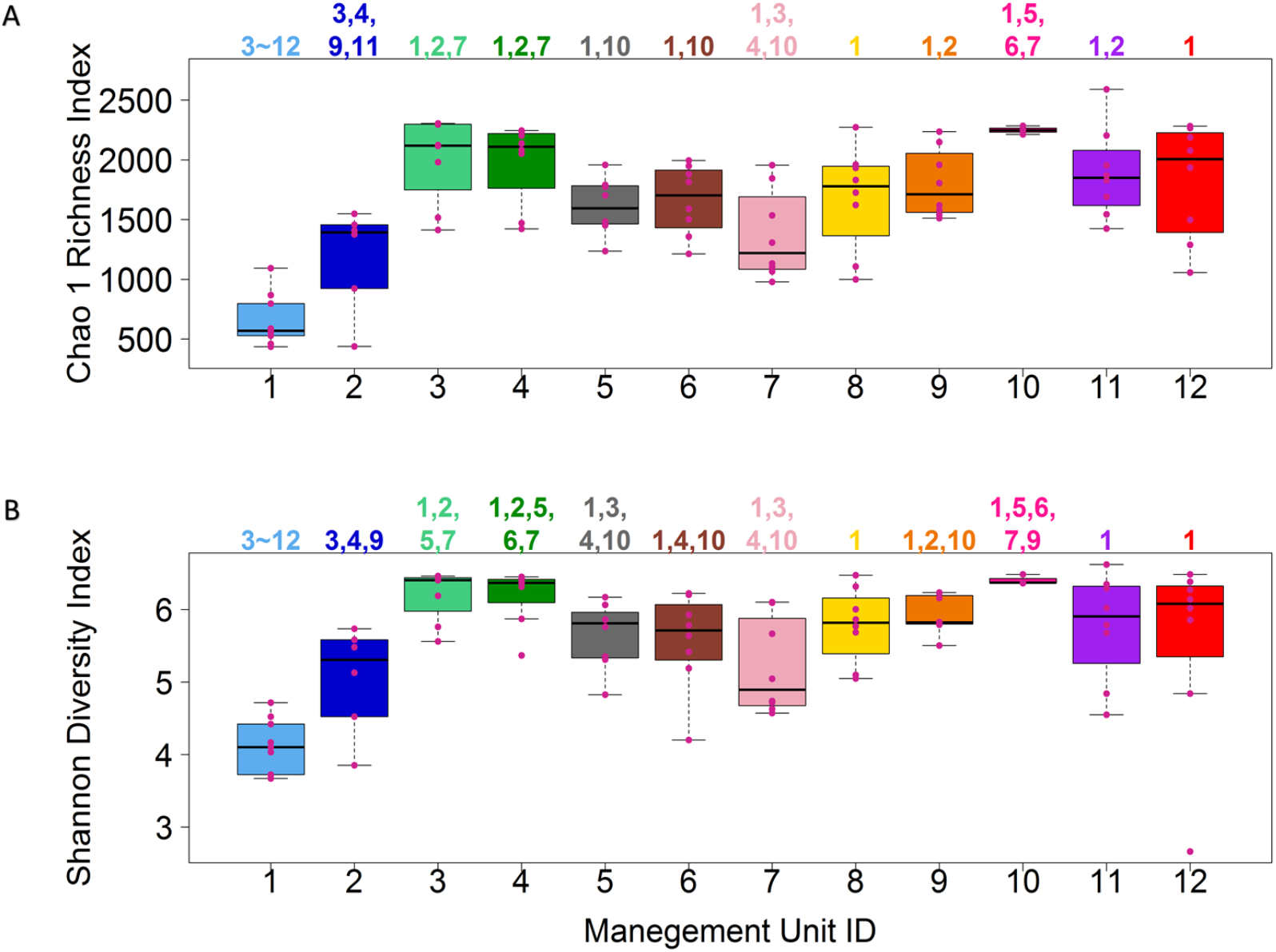
Diversity of fecal bacterial communities of cattle in the management units. Boxplots of Chao 1 richness index (A) and Shannon diversity index (B) of fecal bacterial communities of cattle in the management units (x-axes). Each violet-red dot represents the bacterial communities in a fecal sample and management units are coded by distinct colors. The numbers above each box indicate the management units that are significantly different from the current one with the pairwise Wilcoxon test (P < 0.05).

### Similarity and difference of the fecal bacterial communities between different management units

To further evaluate the similarity and difference of the bacterial communities between different management units, we calculated the *β*-diversity (Bray-Curtis distance) of the samples and visualized the results using non-metric multidimensional scaling (NMDS). As shown in Fig. 4, the hutch calves samples (unit 1) are largely clustered together and clearly separated from the samples from the other management units, with two post weaned heifers (unit 2) samples included, suggesting that the fecal bacterial communities of hutch calves (unit 1) are similar to one another but largely different from those of cattle in the other management units. Interestingly, the samples from the other units form two rather compact distinct clusters, suggesting the samples in each cluster have quite similar bacterial communities. One of these two clusters circled by the black ring contains samples from all the management units except unit 1, in contrast, the other cluster circled by the red ring comprises samples only from units 3, 5, 6, 7, 8, 11 and 12, and not from cattle within two months of calving (non-lactating), regardless of whether they were uni- or multiparous (units 4, 9 and 10), suggesting that such cattle may only have one type of rather uniform bacterial structure. A uniform bacterial structure in such cattle could be related to the type of diet fed to dry-cows.

**FIG 4.**
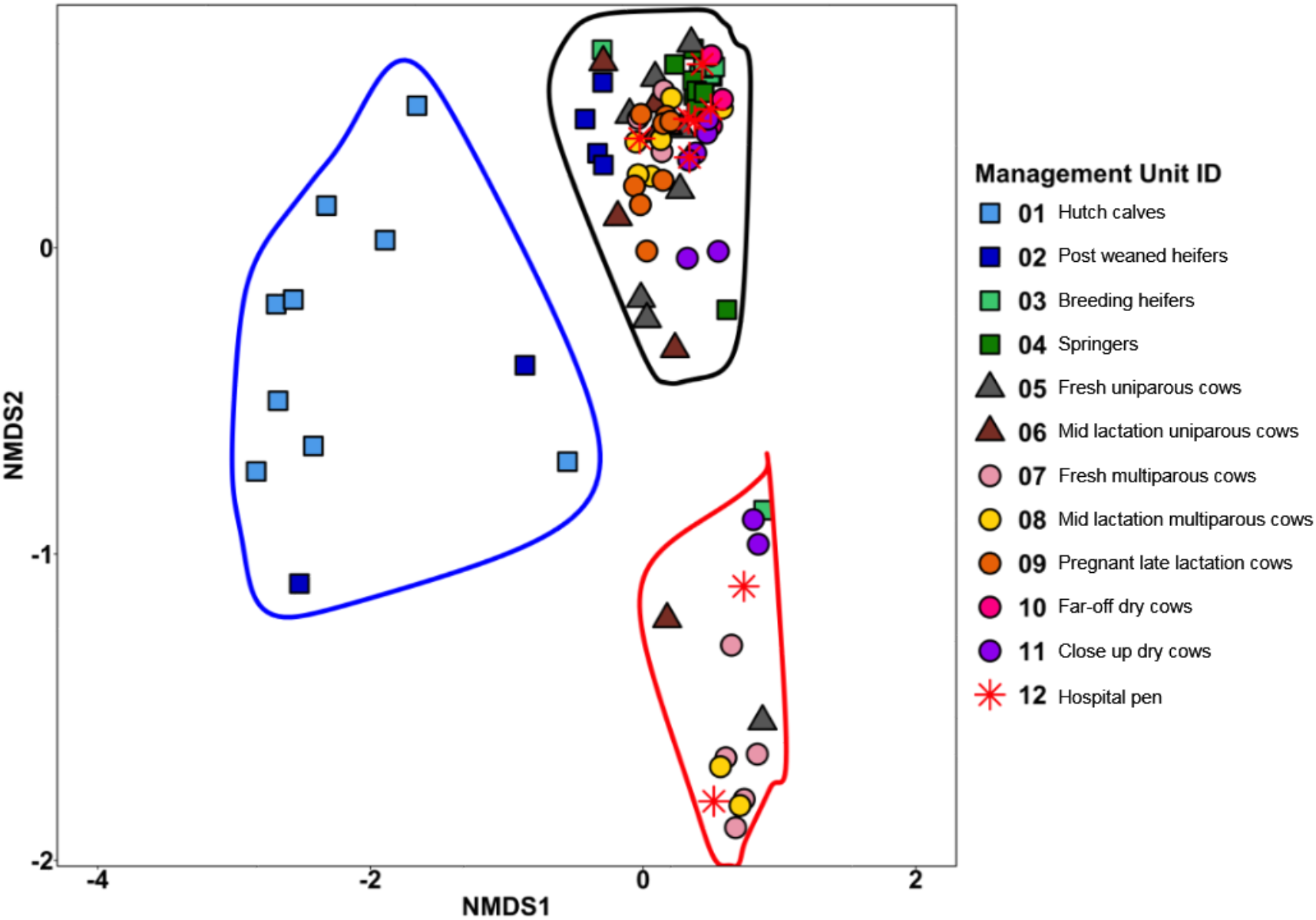
Non-metric multidimensional scaling (NMDS) plot of the Bray-Curtis distance for bacterial communities in different management units. Each point represents a fecal bacterial community and is colored by the management unit from which it was sampled. The communities are grouped into three clusters circled in blue, black and red rings, respectively.

### Shared and unique ASVs in the management units

We examined how the ASVs were shared by the fecal bacterial communities collected from different management units. As shown in Fig. 5, most (4604 ASVs/98.4%) of the 4,681 ASVs were found in the samples from at least three management units. Particularly, 513 ASVs were seen in at least one sample from all the 12 units. On the other hand, 13 and 64 ASVs appeared in the samples from only one and two management units, respectively (Fig. 5), suggesting that they are unique to the relevant units. Out of the 13 unique ASVs, only three ASVs assigned as [o]Bacteroidales (345 reads/0.3%), [f] Prevotellaceae (264 reads/0.2%), and Bacteroides (192 reads/0.17%) had abundances > 0.1% and were all found in hutch calves or unit 1. However, if we slightly loosen the criteria and define “uniqueness” as “an ASV that exists in one unit with a minimum abundance of 0.1% with a total of no more than 3 reads in all other management units”, we found a total of 19 unique ASVs, and all of them were only observed in management unit 1 (Table 2). At the same time, we did not find any such “unique” ASVs in any of the other 11 units.

**FIG 5.**
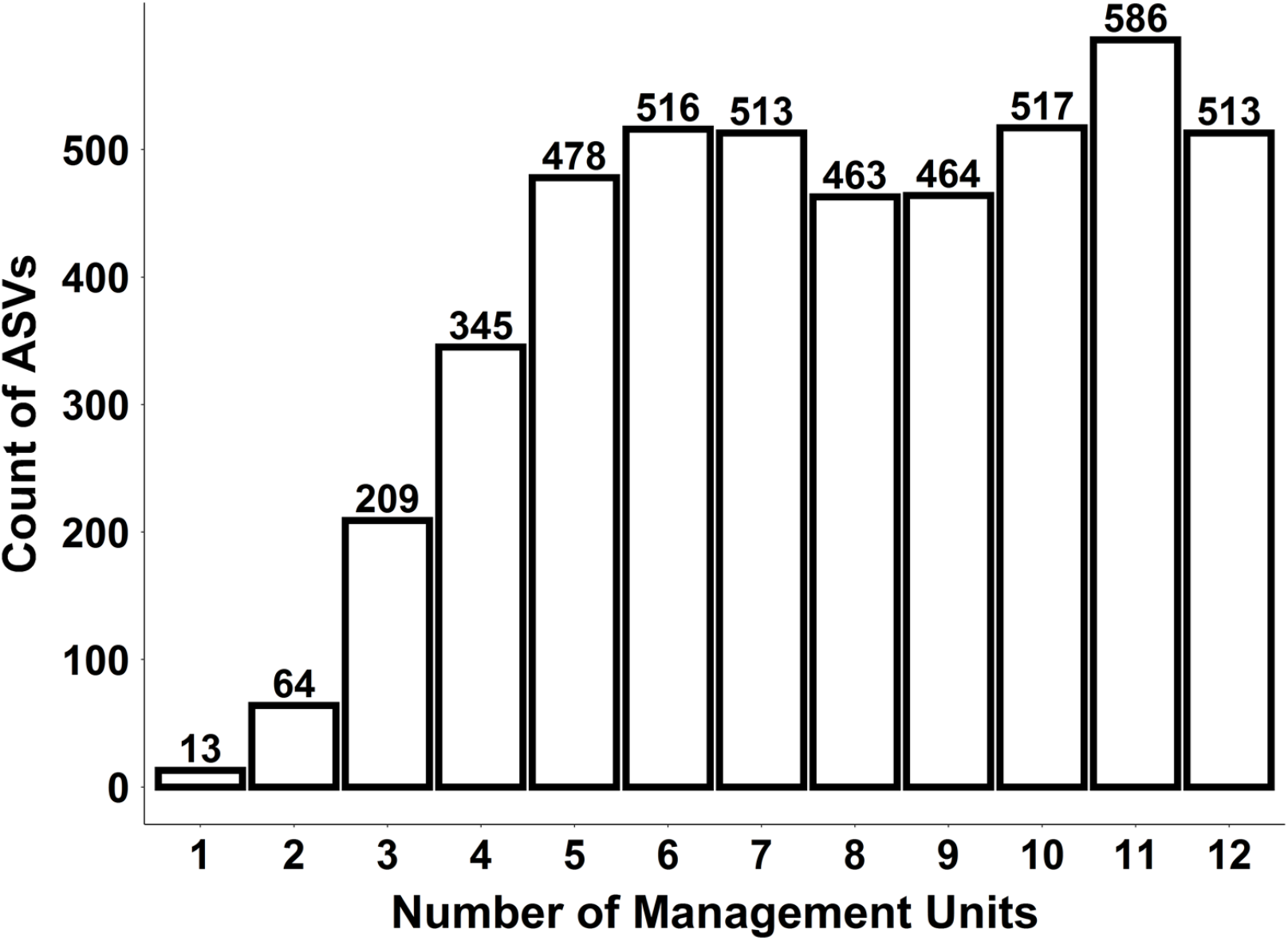
Number of ASVs shared by the fecal samples from different numbers of management units. The x-axis is the number of management units in which the y-axis-indicated number of ASVs were found.

**TABLE 2.** Number of occurrences and taxonomic assignment of ASVs that were exclusively detected in unit 1.

| No. of reads<br>in unit 1 | No. of reads<br>in other units | Phylum | Family | Genus | ASV<br>ID |
| --- | --- | --- | --- | --- | --- |
| 987 | 2 | Bacteroidetes | Prevotellaceae | [f]Prevotellaceae | ASV-260 |
| 525 | 2 | Bacteroidetes | Prevotellaceae | Prevotella_2 | ASV-429 |
| 716 | 2 | Bacteroidetes | Muribaculaceae | [f]Muribaculaceae | ASV-468 |
| 544 | 1 | Bacteroidetes | Prevotellaceae | Prevotella_2 | ASV-471 |
| 455 | 2 | Bacteroidetes | Prevotellaceae | [f]Prevotellaceae | ASV-535 |
| 454 | 2 | Bacteroidetes | Bacteroidaceae | Bacteroides | ASV-576 |
| 345 | 0 | Bacteroidetes | [o]Bacteroidales | [o]Bacteroidales | ASV-640 |
| 346 | 2 | Bacteroidetes | Prevotellaceae | Prevotella_2 | ASV-711 |
| 274 | 2 | Firmicutes | Lachnospiraceae | [f]Lachnospiraceae | ASV-738 |
| 630 | 2 | Firmicutes | Ruminococcaceae | Ruminococcaceae_UCG-014 | ASV-861 |
| 229 | 1 | Bacteroidetes | Muribaculaceae | [f]Muribaculaceae | ASV-1114 |
| 192 | 1 | Bacteroidetes | Bacteroidaceae | Bacteroides | ASV-1131 |
| 264 | 0 | Bacteroidetes | Prevotellaceae | [f]Prevotellaceae | ASV-1150 |
| 192 | 0 | Bacteroidetes | Bacteroidaceae | Bacteroides | ASV-1173 |
| 162 | 2 | Bacteroidetes | Prevotellaceae | [f]Prevotellaceae | ASV-1190 |
| 159 | 1 | Bacteroidetes | Tannerellaceae | Parabacteroides | ASV-1211 |
| 145 | 1 | Firmicutes | Acidaminococcaceae | Phascolarctobacterium | ASV-1222 |
| 209 | 1 | Bacteroidetes | Prevotellaceae | [f]Prevotellaceae | ASV-1286 |
| 129 | 2 | Firmicutes | Erysipelotrichaceae | Erysipelotrichaceae_UCG-004 | ASV-1361 |

### Dynamics of bacterial communities between the management units

From the core bacteria (472 ASVs with a minimal abundance of 0.1% in more than 10% of total samples), we found 212 ASVs displayed significantly different abundances in the samples from the 12 management units (ANOVA, FDR < 0.05). These 212 ASVs can be assigned to 7 phyla, and most of them belonged to the phyla Firmicutes and Bacteroidetes (Fig. 6A). Of the 123 ASVs assigned to phylum Firmicutes, 34 of them were classified as [f]Ruminococcaceae, followed by Ruminococcaceae_UCG-005 (20 ASVs), Ruminococcaceae_UCG-013 (11 ASVs), Ruminococcaceae_UCG-010 (9 ASVs), and a few assigned to other genera (Fig. 6B). Within 75 ASVs assigned to phylum Bacteroidetes, 17 were assigned as Rikenellaceae_RC9_gut_group, while [f]Muribaculaceae and [o]Bacteroidales each accounted for 12 ASVs (Fig. 6C). The relative abundances of these ASVs at three taxonomic levels were shown in Fig. S2 in the supplemental file 2).

**FIG 6.**
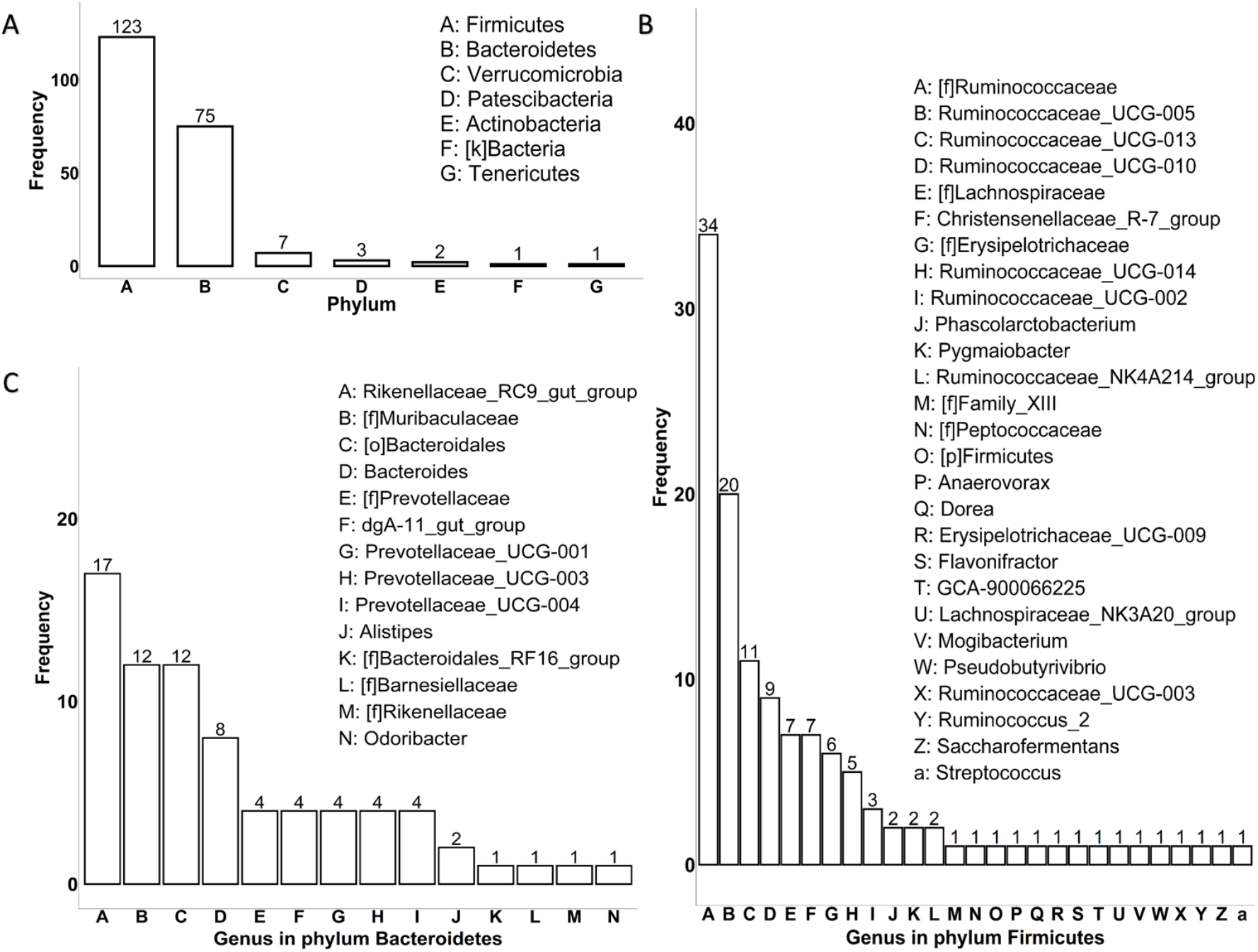
The distribution of the ASVs that display significant differences in abundance in the samples from different management units. (A) The distribution of the number of significantly different ASVs at the phylum level. The distribution of significantly different ASVs (B) in the phylum Firmicutes at the genus level, and (C) in the phylum Bacteroidetes at the genus level.

We further analyzed how each of these 212 ASVs was significantly different in their composition between different pairs of management units (Tukey HSD, p < 0.05). As shown in Fig. 7, unit 1 has 74 ASVs different from unit 3 and 70 ASVs different from unit 4. Many different ASVs were also observed between units 3 and 7 (70 ASVs) and between units 4 and 7 (65 ASVs).

**FIG 7.**
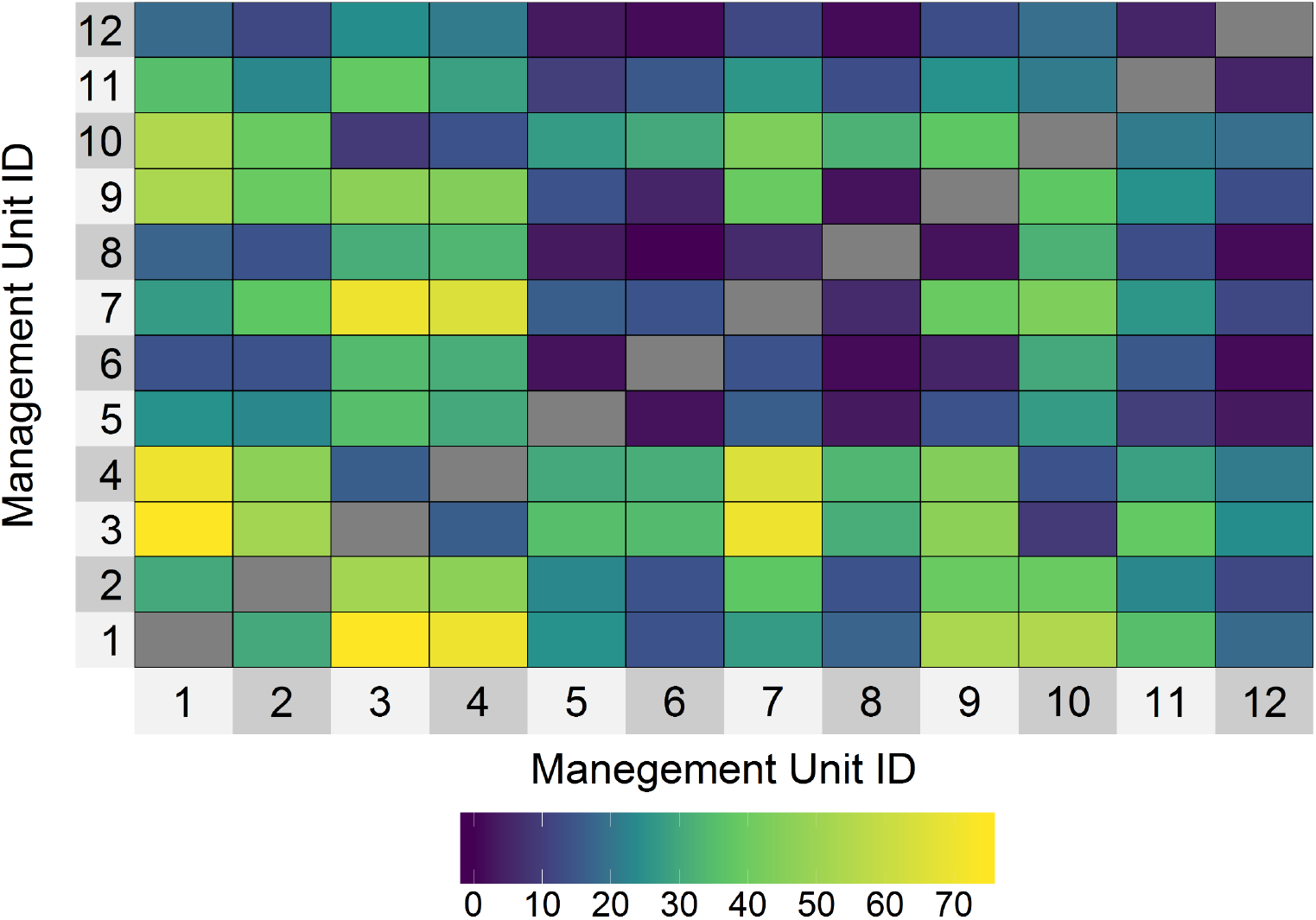
Number of ASVs significantly different between each pair of the management units.

### Synergistic and antagonistic relationships of the ASVs

Bacteria thriving in a community may interact with one another synergistically or antagonistically. To reveal such potential interactions between different bacteria, we calculated the correlations of the core bacteria using the SparCC (Sparse Correlations for Compositional data) algorithm (15). By setting ± 0.7 as the threshold values for strong positive and negative correlations and pseudo p-values < 0.05 as the significant correlations, we found 61 ASVs that had strong correlations with at least one other ASV (Fig. 8; supplemental file 3: SparCC table; supplemental file 4: Taxonomic assignment of 61 strongly correlated ASVs). Most of these strong correlations are positive, while only one pair between ASV-2 (Ruminococcaceae_UCG-005) and ASV-68 (Bacteroides) is negative (SparCC = −0.70). Specifically, ASV-1 (Candidatus_Saccharimonas) has the largest number of strong correlations and highly co-occurred with 36 other ASVs. ASV-36 ([f]Erysipelotrichaceae) has strong correlations with 27 other ASVs. On the other hand, 19 ASVs have a strong correlation with only one other ASV. In Fig. 8, all the strong correlations are shown within the networks. From the 181 strong correlations, 117 are between the ASVs assigned to the same phylum. Out of the 64 strong correlations between the ASVs assigned to different phyla, one is between Bacteroidetes and Firmicutes, the rest all are between Firmicutes and Patescibacteria.

**FIG 8.**
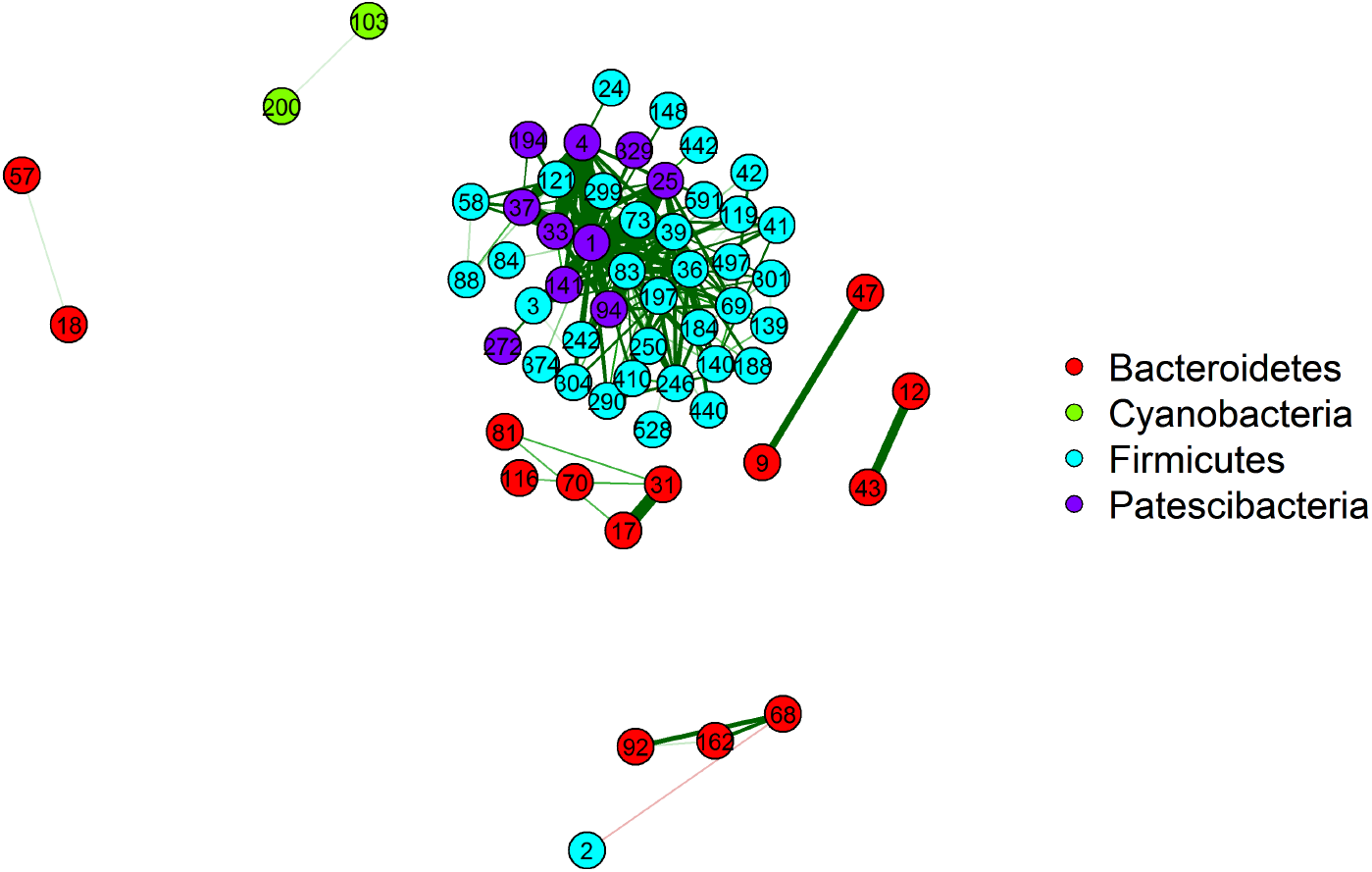
Strong correlations between 61 of the 472 core bacteria. Each node is an ASV and the number within represents the ASV ID. Nodes were colored based on their taxonomic assignment at the phylum level. The edges in green and red indicate positive and negative correlations.

## DISCUSSION

In modern dairy management, it is not uncommon to house animals in different management units based on their ages, nutrition requirements, reproductive status, lactation status, production, and management styles (16). Such a management system optimizes milk production and management but brings challenges profiling and analyzing the bacterial communities in the GIT because the bacterial communities can be influenced by numerous factors including, but not limited to, diet, animal physiology and status, the livestock environment, geographic location, antimicrobials used, and even management practices (5, 11, 17–20).

In this study, we sought to characterize the composition, diversity, and dynamics of the fecal bacteria in 12 management units over the production lifecycle of dairy cattle on a California dairy. We used Illumina Miseq sequencing of the V3/V4 hypervariable regions of the 16S rRNA genes to profile the bacterial communities in 90 fecal samples collected from the 12 management units. Our results suggest significant differences in the 12 management units in terms of richness, evenness, and structure of the bacterial communities.

Overall, consistent with several earlier studies (5, 6, 11, 12, 17), our results reveal Firmicutes and Bacteroidetes were the two most dominant taxa at the phylum level, distantly followed by several other phyla. Specifically, our results showed that, on average, the Firmicutes and Bacteroidetes represented 51.63% and 36.30% of the total communities, respectively. Interestingly, the relative abundance of Bacteroidetes was lower in cows immediately prior to and after calving (within two weeks from calving) in units 7 (18.77%) and 11 (23.25%) while the relative abundances were in the range of 29.48% and 62.25% in other units (Fig. 2B).

For some genus, as Mao et.al demonstrated, Prevotella had lower abundance in samples taken from the small and large intestines, but higher in the forestomach, such as the rumen (6). In our work, the relative abundance of Prevotella was only 0.07%. However, in one of the hutch calves’ (unit 1) samples, we observed Prevotella had a relative abundance of 7.16% (905 reads) while the number of Prevotella reads in the other 89 samples was in the range of 0 to 11. We also found in this sample, Ruminococcus_1 had a relatively high abundance (5.56%, 703 reads) while this genus was in the range of 0 to 115 reads in other samples. So, there may be some synergistic relationship between Prevotella and Ruminococcus_1 and the interaction prevents Prevotella from being degraded or eliminated when going through the GIT. Such an interaction could also be related to the diet of calves in unit 1 being primarily milk compared to all the other units fed a solid diet. Moreover, feeding waste milk from the hospital pen or fresh cow pen to calves is a common practice in the dairy industry and was done on this dairy at the time of sampling. Waste milk commonly contains antibiotic residues of varying concentrations as it is collected from milking cows in the hospital being treated with antibiotics or awaiting clearance during their withdrawal period before rejoining the pens where milk is harvested for human consumption.

It is noticeable that bacterial communities in management unit 1 were significantly different from those in other management units except unit 2 (Figs. 2, 3, 4). These differences confirmed the earlier findings that the enteric microbiota in neonatal calves were different from the adult cows and the bacterial communities underwent a dramatic change during the development in early age (8, 9, 18). Intriguingly, bacteria from the samples in unit 2 displayed similar patterns with the communities from the adult cattle, predominately Firmicutes; but at the same time, they showed no significant difference from those in unit 1. As the cattle in unit 2 were aged from approximately 2 months (70 days average) to 13-month old, the microbiota changes were still ongoing towards the bacterial profiles in adults at this transition period. Using the same set of samples, we have previously found *E. coli* from unit 1 (hutch calves) exhibited a wider spectrum of resistance to antimicrobial drugs compared to bacteria from other units (16). Currently, it largely remains undetermined with respect to the roles of bacterial communities on antimicrobial resistance of specific bacterial species, however, in future studies, it will be interesting to assess such roles, for example, resistomes on phenotypes of antimicrobial resistance.

For the samples from non-neonatal cattle (from unit 2 to unit 12), we did not find any evidence supporting that the bacterial communities could be clustered based on the management unit membership. This demonstrated that the management unit itself is not a determinate force for the structure of the bacterial communities. Instead, as shown in Fig. 4, we did observe distinct patterns where most of our samples from management unit 2 to management unit 12 formed two major clusters, implying they had two major different structures. As we looked deeper, we found that at the phylum level, the relative abundance of Patescibacteria became very different between the two clusters (Fig. S3 in the supplemental file 2). In the red circled cluster, the mean and median of Patescibacteria relative abundance are 20.87% and 19.66% while those values in the black circled cluster were only 1.90% and 1.07%, respectively. What caused the differences in Patescibacteria in various samples still needs future research, but the potential interactions between some taxa classified as Firmicutes and Patescibacteria seem to prevent Patescibacteria from decreasing. It has been reported that the rumen bacteria in the 1^st^ and 2^nd^ lactations, which overlapped with some of our management units, are dynamic yet similar, and the samples from the two lactations cannot be separated by the lactation cycle in PCA visualization (7). This demonstrated that the bacterial structure might undergo further shaping going through the GIT (their rumen samples versus our fecal samples) or the formed patterns were just a case-sensitive phenomenon.

Pitta et.al reported that significant bacterial population change was observed during the transition from 21 days prior to calving to 21 days after calving in uniparous and multiparous cows, respectively (21). This shift period corresponded approximately to our management units 4 and 5 for uniparous cows and units 10 and 11 for multiparous cows. However, we did not see this significant shift as demonstrated earlier (21). The only significant difference we observed was Shannon diversity between units 4 and 5. This could be due to the different sites of the GIT from which the samples were taken (their rumen samples versus our fecal samples). It is also possible that this was due to the four management units in our study not being strictly narrowed to 21 days prior and post-calving. As the dairy farm environment is dynamic, it is not surprising that most taxa were shared by cattle housed in different management units.

Measuring potential bacterial interactions was challenging and barely reported in the dairy cattle studies. The classic correlation methods had their limitations when applied to genomic data such as 16S rRNA sequencing data which are sparse and compositional. SparCC as a method that negates the negative correlation bias of compositional data (22) and identifies true association missed by others (23), was used here as a way of evaluating bacterial interactions. In general, we identified 180 strong positive correlations and 1 strong negative correlation between 61 ASVs (Fig. 8). These co-occurrent ASVs may tend to share the same habitats and perform similar functions as most of these co-occurrent patterns spotted between ASVs were in the same phyla (Fig. 8).

In conclusion, in this study, we profiled the structure and dynamics of bacterial communities from cattle in 12 independent yet inter-connected management units on a modern California dairy. To the best of our knowledge, this is the first study that describes the bacterial communities across all management units and clearly depicts the microbial communities to each of the clearly defined management units. We demonstrated the dynamics of change across production systems and confirmed that microbial ecology underwent dramatic changes in the early days of life, as evidenced by the significantly different bacteria in hutch calves from other adult cattle. Moreover, we identified 19 ASVs that were detected only in hutch calves in management unit 1 but were absent in all the other cattle in other units. These ASVs might play crucial roles in the early establishment and development of the GIT. We also dissected the details of the bacterial dynamics across the management units and reported potential bacterial interactions.

## MATERIALS AND METHODS

### Sample collection and study herd

Using standard veterinary protocols, a total of 90 fecal samples were collected from the rectum of 90 dairy cattle in 12 management units on a single day in June 2016 from a dairy herd in the Central Valley of California, USA. Sampling was approved by the Institutional Animal Care and Use Committee (IACUC) of University of California Davis (protocol number 18941). The schematic diagram of the management units is shown in Fig. 1 with the description of the management units and cattle in Table 1. Other information such as the sampling population in each management unit can be found at Li et.al (16).

### Illumina MiSeq Sequencing

Two steps of Polymerase Chain Reaction (PCR) procedures were used to generate amplicons from the 16S RNA genes for sequencing. The first-round PCR was to target V3/V4 regions of 16S rRNA genes with the forward primer: 5’-CCTACGGGNGGCWGCAG and the reverse primer: 5’-GACTACHVGGGTATCTAATCC. This step was done with the KAPA Biosciences HiFi PCR kit and additional BSA. The protocol consists of initial denaturation at 95 °C for 3 min, followed by 25 cycles of denaturation (90 °C for 30 s, 55 °C for 30 s, and 72 °C for 30 s), and final elongation at 72 °C for 5 min. The PCR products were cleaned up with Ampure XP beads. The second-round PCR was performed with Nextera XT index Primers and sequencing Adaptors with the following setting: initial denaturation at 95 °C for 3 min, followed by 8 cycles of denaturation (90 °C for 30 s, 55 °C for 30 s, and 72 °C for 30 s), and a final elongation at 72 °C for 5 min. The PCR products were cleaned up with Ampure XP beads and paired-end sequenced (2 x 300bp) on an Illumina MiSeq platform at the University of North Carolina at Charlotte.

### ASV Table Construction

Primers with raw sequences were removed by Cutadapt (24). We performed quality control using DADA2’s “filterAndTrim” function with “trancLen” equal to 200bp for forward reads and 150bp for reverse reads based on quality profiles. Technical error rate learning was performed with all the sequences in the samples. Sample inference was performed by the “dada” function with the setting optional parameter “pool=TRUE.” Paired-end reads merger in DADA2 resulted in approximately 50% loss of sequences, thus only forward reads were used in this study. ASV table and chimera removal were implemented with default parameters. We used the “assignTaxonomy” function that provided a native implementation of the RDP Classifier (25) with minimum bootstrap confidence of 80 to assign taxonomy from the phylum level to the genus level to each ASV. SILVA database release 132 (26) was used as the reference database.

### Statistical Analysis

Rarefying was performed by the “single_rarefaction” function with the minimum number of reads of all samples, which is equal to 12,637 in Qiime (27). We used alpha diversity, including Chao 1 index that evaluates richness and Shannon index that evaluates diversity to measure the within-sample diversity. Differences in Chao 1 index and Shannon index in different units were assessed by Kruskal Wallis test with the Benjamini-Hochberg (BH) correction for multi-comparison, and pairwise management units comparisons were performed by pairwise Wilcoxon test. The Bray-Curtis distance matrix was employed to perform beta diversity analysis. Visualization was done by non-metric multidimensional scaling plot. SparCC correlations were calculated by FastSpar (28), a C++ implementation of SparCC algorithm (100 bootstrap samples were generated for pseudo p-value calculation). R software (29) and R packages ggplot2 (30), vegan (31), superheat (32), ggpubr (33), and qgraph (34), were used for calculation and visualization.

### Accession numbers

All DNA sequences have been deposited in NCBI’s Sequence Read Archive (SRA) with the BioProject access number PRJNA607283.

## SUPPLEMENTAL MATERIAL

SUPPLEMENTAL FILE 1: Normalized ASV table, TXT file

SUPPLEMENTAL FILE 2: Supplemental figures, PDF file

SUPPLEMENTAL FILE 3: SparCC table, TXT file

SUPPLEMENTAL FILE 4: Taxonomic assignment of strongly correlated ASVs, TXT file

## ACKNOWLEDGMENTS

This study was partially supported by the National Institute of Food and Agriculture (NIFA)’s Exploratory Research program (2015-67030-23892), the NIFA’s Dairy Herd Health and Food Safety Formula Funds (CALV-DHHFS-0053) from the USDA, and National Institutes of Health (R01GM106013).

